# Anthropogenic disturbance inverts microclimate stratification in tropical forests

**DOI:** 10.64898/2026.06.11.731463

**Authors:** Cássio Alencar Nunes, Erika Berenguer, Rodrigo O. do Nascimento, Raquel Gasparini Martins, Oliver C. Metcalf, Alexander C. Lees, Marielle N. Smith, Joice Ferreira, Ilya Mclean, Jos Barlow

**Affiliations:** Lancaster Environment Centre, Lancaster University, Lancaster, United Kingdom; Environmental Change Institute, University of Oxford, Oxford, United Kingdom; Embrapa Amazônia Oriental, Belém, Brazil; Department of Natural Sciences, Manchester Metropolitan University, Manchester, United Kingdom; School of Environmental and Natural Sciences, Bangor University, Bangor, United Kingdom; Environment and Sustainability Institute, University of Exeter, Penryn, United Kingdom

**Keywords:** Amazonia, Fire, Forest structure, Microclimate buffering, Temperature, Understorey

## Abstract

Tropical rainforests generate and maintain their own microclimate regimes and the resultant cooler, more humid and stable environments foster the hyperdiversity typical of these ecosystems. Although the temporal and spatial (horizontal) distributions of microclimates have been relatively well studied, the vertical dimension has received less attention, and little is known about how forest disturbance affects the vertical stratification of microclimates in tropical forests. In this study, we examine how the vertical distribution of temperatures varies between undisturbed and burned Amazonian forests. We installed five vertical transects with temperature dataloggers distributed at 7 different heights to collect data over multiple days during the end of the dry season. We investigated how anthropogenic disturbance (fire) mediates the vertical stratification of microclimate and whether microclimate buffering (i.e, the difference between understorey and canopy temperatures) varies according to the forest structure. We showed that anthropogenic disturbance can cause an inversion in the vertical stratification of microclimates, with burned forests having hotter temperatures (up to 2 °C) in the understorey than in the canopy during the day – the opposite of what is found in undisturbed forests (typically 3 °C cooler). During the night, while understorey and canopy temperatures are similar in undisturbed forests, we found that, in burned forests, understorey temperatures were up to 2 °C cooler than in the canopy. Microclimate buffering by day was best explained by aboveground carbon stocks, with higher temperature buffering in more carbon rich forests. Our study shows that anthropogenic disturbance alters the vertical stratification of temperatures in Amazonian forests, leading to significant temporal changes along the diel cycle. Future research should focus on understanding these changes across a wider range of disturbance regimes, and explore the consequences for biodiversity and ecosystem functions from the canopy to the forest floor.

## 1 INTRODUCTION

Tropical forests are the most biodiverse terrestrial ecosystem on Earth (Pillay et al. 2022) and provide ecosystem services ranging from local pollination to contributing to global climate regulation (Bonan 2008). The hyperdiversity in tropical forests is partially supported by their ability to buffer the climate above the canopy, creating a cooler, more humid and more stable microclimate – understorey temperatures in tropical forests are, on average, 1.6 °C cooler than in the open air above the canopy (Ismaeel et al. 2024). This microclimatic buffering influences species distributions and interactions (De Frenne et al. 2021, Kemppinen et al. 2024) and affects ecosystem functioning [e.g., wood decomposition (do Nascimento et al. 2026), sapling recruitment and nutrient cycling (Jucker et al. 2020)]. In addition, it can reduce the risk of extinction from climate change by creating microrefugia within the forest (Suggitt et al. 2018). Finally, microclimate buffering is of particular relevance given the growing prevalence of tropical forest fires as hotter and drier microclimates increase the chances of fire occurrence (Uhl et al. 1988, Ray et al. 2005).

Tropical forest microclimates vary temporally and spatially. Temporal variations of microclimate are relatively well studied. Microclimate shows clear diel trends, with temperature peaking in forests at around mid-afternoon (Ma et al. 2025). Spatial variation of microclimate can be horizontal, *i.e.*, the distribution of understorey microclimates within forests (and between forests and other ecosystems) and vertical, *i.e.*, the distribution of microclimates across the forest strata. The horizontal distribution of microclimates in tropical forests has received considerable recent attention and it is known to be driven by a combination of topography, water balance (through evapotranspiration), wind and vegetation structure (De Frenne et al. 2021). The magnitude of difference between understorey and open air temperature will be larger when macroclimatic temperatures are higher (increased buffering), water availability for trees is higher (increased evapotranspiration), the distance from forest edge is higher (lower edge effects) and vegetation structure is well preserved (increased shading) (Ewers and Banks-Leite 2013, Jucker et al. 2018, De Frenne et al. 2021, Ismaeel et al. 2024, Ma et al. 2025). Forest disturbances affect many of these drivers, and any events that open up the canopy – such as selective logging or fire (Berenguer et al. 2014) – result in warmer and drier understorey microclimates (Santos et al. 2024).

Vertical stratification of microclimate in tropical forests typically leads to higher temperatures by day near the canopy which decrease towards the forest floor, while at night temperatures are more stable across the vertical profile (De Frenne et al. 2021). This pattern is caused by the stratification of vegetation that intercepts light and changes air movement, leading to layers of microclimates from canopy to forest floor depending on the distribution of leaves (De Frenne et al. 2021). However, in contrast to our increasingly strong understanding of how forest disturbance impacts the horizontal distribution of microclimates (Jucker et al. 2018, Santos et al. 2024), little is known about how disturbance affects the vertical stratification of microclimates. Forest disturbance affects canopy openness and the leaf layers in the forest strata, increasing the penetrance of light into the forest understorey (Fauset et al. 2017). Although a reduction in light interception will likely cause an increase in temperatures near the forest floor (as evidenced for the horizontal distribution), we do not know exactly how it changes temperature in different layers from the canopy to the forest floor. In addition, we lack information on how disturbance-induced changes in vegetation structure affect the temporal patterns of the microclimate across different heights in the forest.

Here, we examine how the vertical distribution of microclimate changes between undisturbed and burned forests in Amazonia. To answer this, we installed 29 sensors, each collecting data between 8 and 16 days during the dry season of 2024. Specifically, we ask: (**Q1**) Does the vertical stratification pattern of microclimate change between undisturbed and burned forests? We looked at the variation in average temperatures with height and throughout the day; (**Q2**)

How does anthropogenic disturbance change the microclimate buffering at different heights? And (**Q3**) is the microclimate buffering capacity/loss explained by the loss of forest structure?

## 2 METHODS

### 2.1 Study area

The study was conducted in the eastern Brazilian Amazon, within the Tapajós National Forest protected area (3°31’01” S, 55°04’23” W) in the municipality of Belterra, State of Pará. The climate in the region is classified as hot and humid, with a mean annual temperature of 25 °C and mean annual precipitation of 1920 mm (Parrotta et al. 1995). There is a wet season typically extending from December to July and a dry season from August to November, when the monthly precipitation is below 100 mm.

### 2.2 Sampling design

Data was collected from five plots (250 × 10 m) located in *terra firme* (unflooded) primary forests between 24^th^ November and 11^th^ December 2024, during the transition from the dry to the wet season. Plots were distributed along an anthropogenic disturbance gradient going from undisturbed (1 plot; UF, Fig S1a), burned once (2 plots, burned in 2015; B1, Fig S1b and S1c) and burned twice (2 plots, burned in 2015 and 2023; B2, Fig S1d and S1e) primary forests. Plots were between 2 and 45 km apart.

In each plot, we established a vertical transect from the ground to the canopy and installed SurveyTag (https://surveytag.co.uk/) temperature dataloggers. This datalogger measures the air temperature using an ultra-fine wire thermocouple sensor which minimises the effect of solar radiation on air temperature readings. We used a nylon rope to attach the dataloggers at known heights from the ground (Fig. S2), using trees higher than 30 m (emergent above the canopy) in each site to provide us with the most complete overview of vertical stratification in each plot. The number of dataloggers varied between five and seven in the plots due to the variable canopy height (Table 1), but the full set of heights from the ground was: 0 m, 0.5 m, 1 m, 5 m, 9 m, 15 m and 30 m. Dataloggers registered the air temperature every minute during the entire day. Data was collected for 16 days continuously in the UF plot, for eleven days on one B1 and one B2 plots and for eight days in one B1 and one B2 (Table 1).

**Table 1.** Sampling heights and dates in each sampled vertical transect in the Amazon.

| Plot | Heights | Start date | End date | N of days |
| --- | --- | --- | --- | --- |
| UF | 7 (0, 0.5, 1, 5, 9, 15 and 30 m) | 25/11/2024 | 10/12/2024 | 16 |
| B1-a | 6 (0, 0.5, 1, 5, 15 and 30 m) | 24/11/2024 | 01/12/2024 | 8 |
| B1-b | 6 (0, 0.5, 1, 5, 9 and 15 m) | 01/12/2024 | 11/12/2024 | 11 |
| B2-a | 5 (0, 0.5, 1, 5, and 15 m) | 01/12/2024 | 11/12/2024 | 11 |
| B2-b | 5 (0, 0.5, 1, 5, and 15 m) | 24/11/2024 | 01/12/2024 | 8 |

### 2.3 Forest structure

In each plot we measured and identified all trees ≥ 10 cm diameter at breast height (DBH). We also established five subplots (20 × 5 m), in which we measured and identified all trees ≥ 2 cm DBH < 10 cm DBH. Height of each tree was visually estimated. We then used Chave’s (Chave et al. 2014) biomass equations that calculate tree biomass based on DBH, height and wood density. We obtained species-level wood density from the Global Wood Density Database (Zanne et al. 2009) filtered for South American tropical regions. When species-specific wood density values were not available, we used those from the next taxonomic level (*e.g.*, genus-level data). To calculate plot-level biomass, we summed the biomass of every individual. We then extrapolated the value to the hectare of individuals ≥ 10 cm DBH by multiplying by four, while for individuals < 10 cm DBH, we multiplied by 20. Carbon was assumed to account for 50% of biomass.

### 2.4 Data analysis

To investigate how anthropogenic disturbance (*i.e.*, fire) changed the vertical stratification pattern along the day (**Q1**), we modelled the diel cycle of air temperature per height and per plot. We first calculated the mean temperature per minute of the day based on the multiple days that were sampled. We then used Generalised Additive Models (GAM) to model the temperature along the day and used cubic splines to smooth the variation of the mean temperature per minute. We displayed the modelled diel cycle using the ‘*geom_smooth*’ function of the “ggplot2” R package (Wickham 2016, R Core Team 2024). To investigate how anthropogenic disturbance affected the variation of microclimate with height (**Q1**), we calculated the daily mean temperature per height per plot. We divided the data into two slots of time to represent day and night times by using 12:00-04:00 pm and 12:00-04:00 am, respectively. These times represent temperatures of both periods of the diel cycle without including transitioning temperatures that might cause over/underestimation of the mean.

To address **Q2** about how anthropogenic disturbance affects the microclimate buffering (i.e., Δ Temperature between the canopy and other heights) in Amazonian forests, we first calculated the buffering per minute of each plot. We first calculated the mean temperature per minute, per height and per plot. We then subtracted the canopy temperature (15 or 30 m height, depending on the plot; see Table 1) from the air temperature measured at the four heights closest to the ground (0, 0.5, 1 and 5 m – understorey microclimate). This Δ Temperature will therefore be negative when there is microclimate buffering, i.e., when understorey temperature is lower than canopy temperature. We then used GAMs to model the Δ Temperature along the diel cycle, using cubic splines as the smoothing factor.

Finally, to understand if microclimate buffering was explained by changes in forest structure (**Q3**), we calculated the average Δ Temperature during the periods of day and night (according to our classification between 12:00-04:00 am/pm) and used it as a response variable in a linear regression with aboveground carbon stock (Mg/ha) as an explanatory variable in R (R Core Team 2024).

## 3 RESULTS

### 3.1 Temporal changes in temperature across different heights

Mean temperature varied along the diel cycle, with lowest temperatures around 6 am and highest temperatures around 3 pm in all forests (Fig. 1). In undisturbed forests, differences in temperature among different heights were minimal between 6 pm and 6 am. In these forests, vertical stratification of temperatures started to appear from sunrise around 6 am, being more apparent between midday and 6 pm, with understorey temperatures lower than canopy ones (Fig. 1). In burned forests, this diel pattern changed: differences between temperatures from different heights were smaller in forests that burned once, while the forests that had burned twice either showed almost no difference between temperatures from different heights or showed an inverted pattern where understorey temperatures were higher than in the canopy (Fig. 1).

**Figure 1.**
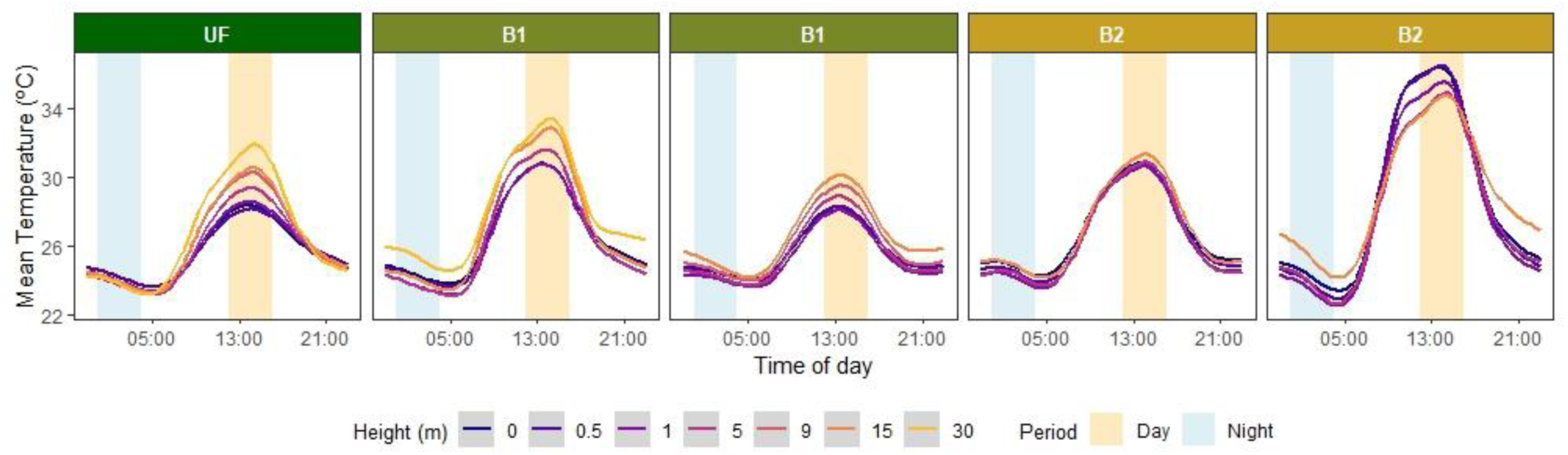
Diel cycle of the vertical stratification of microclimate in undisturbed and burned Amazonian forests. We calculated the mean temperature per minute, per plot and per height band. All heights were measured from the forest floor. We used Generalised Additive Models to model the temperature along the diel cycle. Lines represent model results. Day period is equivalent to the period between 12 pm and 4 pm and night period is equivalent to the period between 12 am and 4 am. UF: undisturbed forest, B1: forest that burned once in 2015. B2: forest that burned in 2015 and 2023.

### 3.2 Changes in average temperatures at each height

When we looked at the average temperatures at each height during the day (12-4 pm), those of undisturbed forest were more elevated in the canopy with values from near the forest floor (0.5 m) being the coolest (Fig. 2). At 0.5 m in the undisturbed forest, temperatures were on average 3.25 °C lower than in the canopy (30 m). This is in stark contrast to the results from one of the forests that burned twice, where temperatures at 0.5 m were on average 1.87 °C higher than in the canopy (Fig. 2). There were also different patterns during the night (12-4 am): while the average undisturbed forest temperatures were all similar at different heights, all burned forests presented cooler temperatures below the canopy (Fig. 2).

**Figure 2.**
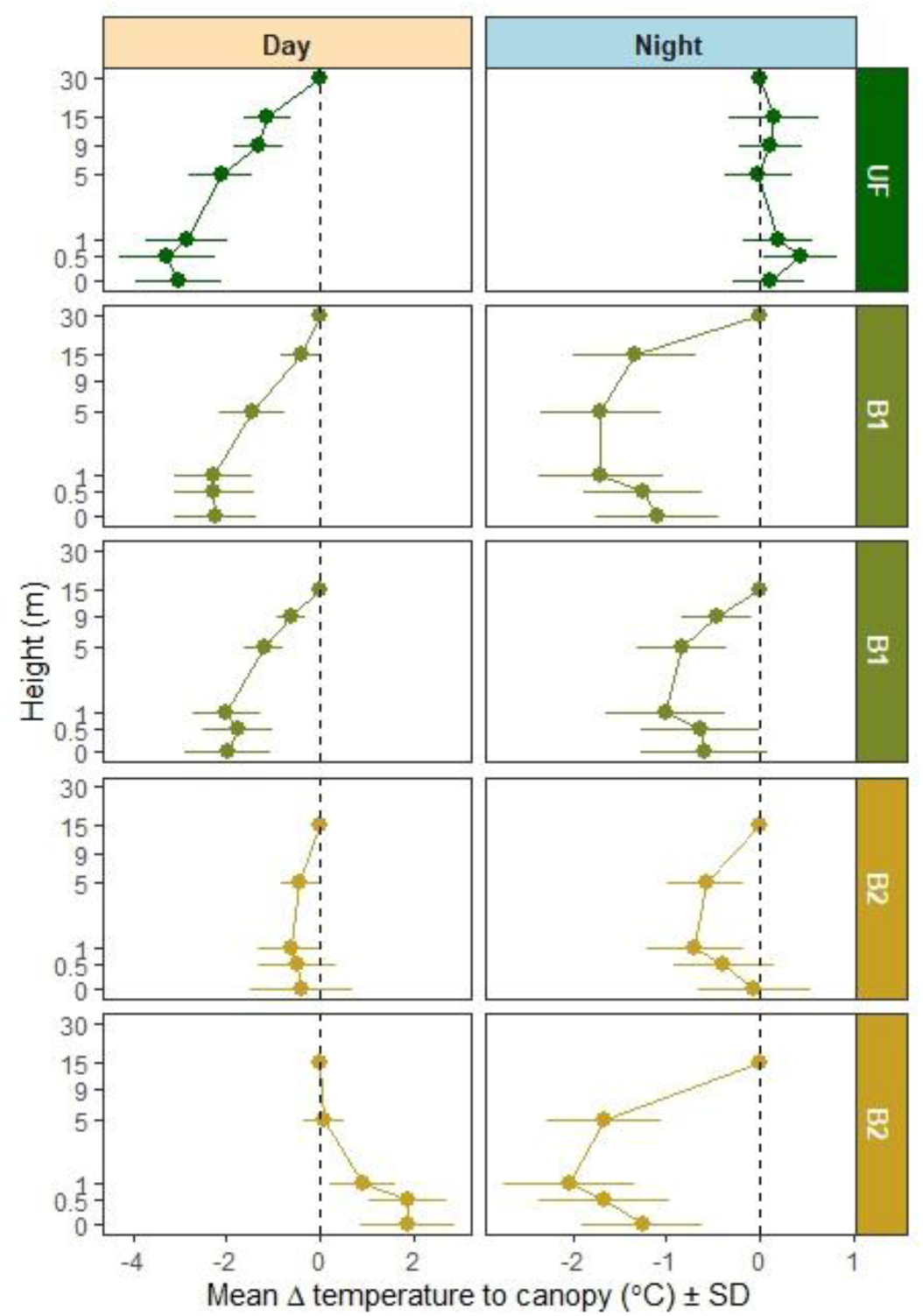
Patterns of vertical microclimate stratification in undisturbed and burned Amazonian forests. As the mean daily temperature was calculated over different days for different plots, we standardised the values by subtracting all values by the value of the highest height (canopy) of each plot. Day period is equivalent to the period between 12 pm and 4 pm and night period is equivalent to the period between 12 am and 4 am. The values in Y axis were log-transformed to facilitate the visualisation of the patterns. UF: undisturbed forest, B1: forest that burned once in 2015. B2: forest that burned in 2015 and 2023.

### 3.3 Changes in microclimate buffering

The microclimate buffering (i.e., Δ Temperature between the canopy and other heights) diel cycle followed the same patterns in the four understorey heights (0, 0.5, 1 and 5 m), although with different magnitudes (Fig. 3). In addition, the inversion of the pattern observed in burned forests when looking at average temperatures was also observed for the microclimate buffering, although again with varying magnitudes at different heights (Fig. 3).

**Figure 3.**
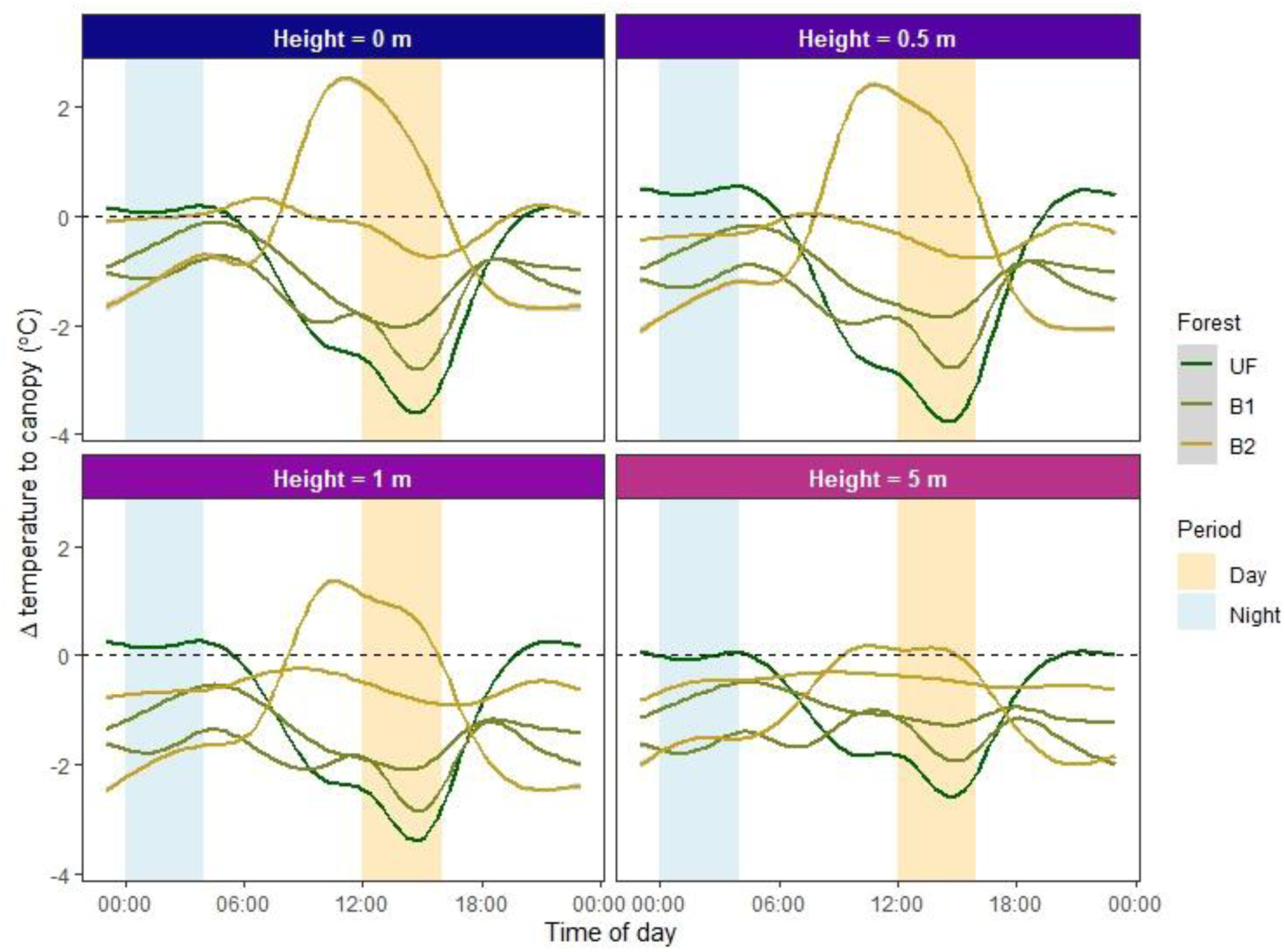
Diel cycle of microclimate buffering in undisturbed and burned Amazonian forests. Temperature of canopy height (30 or 15 m depending on the plot – see Table 1) was subtracted from each of the four heights shown. The mean Δ Temperature was calculated for each minute and modelled using Generalised Additive Models. Day period is equivalent to the period between 12 pm and 4 pm and night period is equivalent to the period between 12 am and 4 am. UF: undisturbed forest, B1: forest that burned once in 2015. B2: forest that burned in 2015 and 2023.

Finally, microclimate buffering during the day was explained by aboveground carbon stocks (Fig. 4). The relationship between aboveground carbon stock and Δ Temperature during the day was negative, meaning that the buffering is higher in forests with higher carbon stock (Fig. 4). During the night, there was no significant relationship between microclimate buffering and aboveground carbon (Fig. 4).

**Figure 4.**
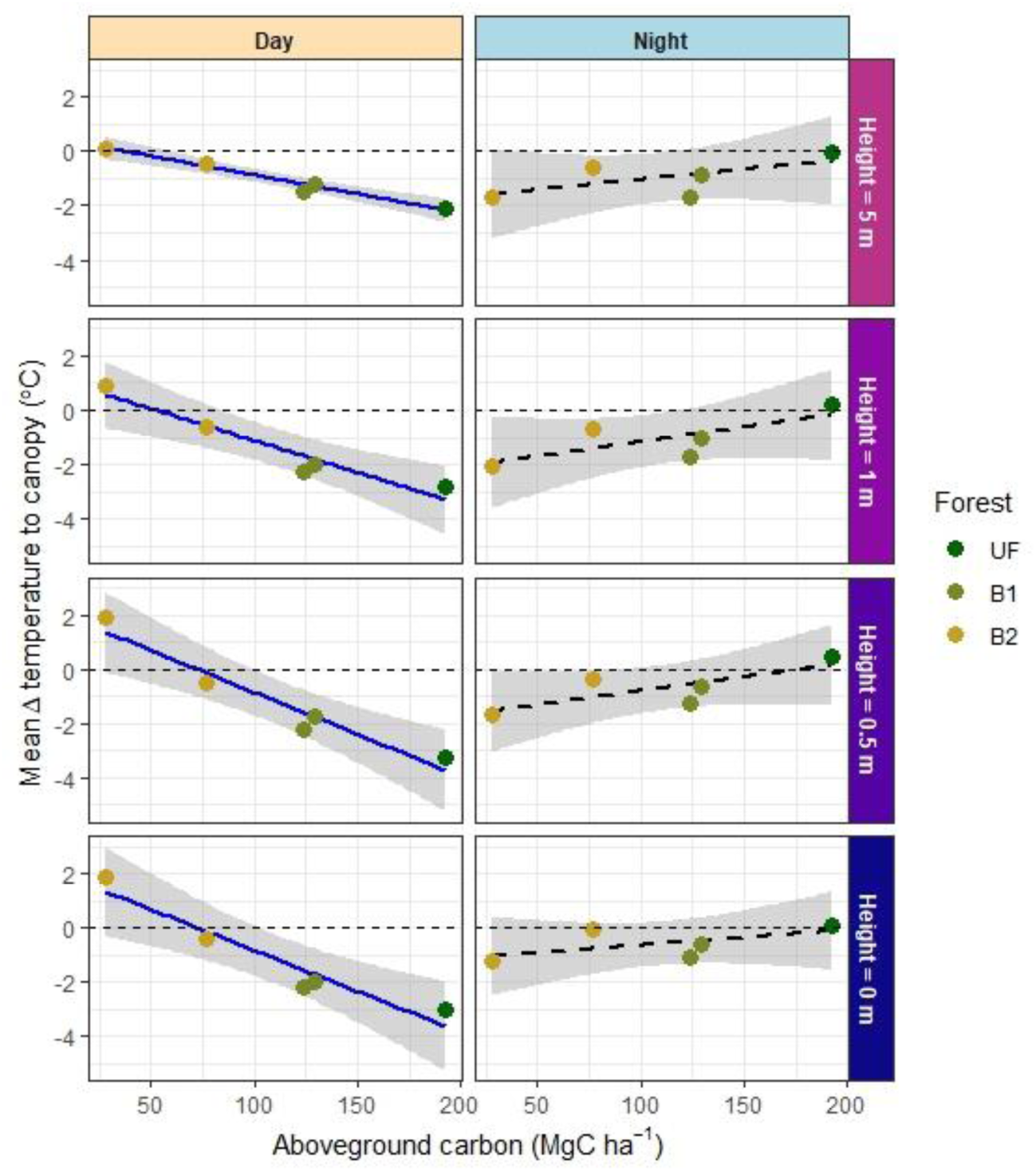
Relationship between microclimate buffering and forest structure in Amazonian forests. The aboveground carbon stock was used as a proxy for forest structure and modelled as an explanatory variable of the mean Δ Temperature to the canopy for the periods of day and night. Day period is equivalent to the period between 12 pm and 4 pm and night period is equivalent to the period between 12 am and 4 am. Solid blue curve represents a significant relationship between mean Δ Temperature and aboveground carbon stock, while dashed black curve represents non-significant relationship. UF: undisturbed forest, B1: forest that burned once in 2015. B2: forest that burned in 2015 and 2023.

## 4 DISCUSSION

Our results show that anthropogenic-induced disturbance can cause an inversion in the vertical stratification of microclimate in Amazonian forests. During the day, undisturbed forests had near-ground temperatures 3.25 °C lower than the canopy on average, while the temperature in forests that burned twice was up to ∼2 °C higher at the forest floor than in the canopy. While undisturbed forest exhibited uniform nocturnal vertical temperature profiles, burned forests had understorey temperatures up to 2 °C cooler than in the canopy on average.

Although this study only presents results on temperature, the findings could have important implications for forest specialists that are adapted to the cooler and more humid environments below the canopy. Research has shown that Amazonian understorey insectivorous birds strongly select dim and cooler microclimates within the forest (Jirinec et al. 2022) and are unlikely to thrive under or adapt to the conditions we found in burned forests. In addition, a comprehensive analysis on Andean-Amazonian insects showed that they have very limited thermal tolerance (Holzmann et al. 2026). The change in the vertical stratification means that not only the fauna that inhabits the forest floor (or near the forest floor) is affected, but that the vertical partitioning of niches is compromised, potentially reducing foraging, roosting or nesting space and disrupting interactions and competition (Nakamura et al. 2017, Kemp et al. 2023). Increased temperature ranges along the vertical profile can also affect the establishment and survival of epiphytes (Gehrig-Downie et al. 2011) that create microhabitats and serve as important resources for fauna (Scheffers et al. 2014). In summary, our study shows that human-induced modification of tropical forests alters not only the horizontal pattern of microclimate but also the vertical stratification of temperatures, including with significant temporal changes along the diel cycle.

Our results also suggest that loss of microclimate buffering and breakdown in vertical stratification of temperatures in human-modified Amazonian forests can be predicted by simplified proxies of forest structure such as aboveground carbon stock. Aboveground carbon stock is higher when trees are taller and larger and when they have higher wood density (Berenguer et al. 2015). Although this metric simplifies many aspects of the vertical stratification of vegetation in the forest (e.g., leaf area index, canopy height, canopy openness) into a single value, it shows a clear response to fire disturbance in the Amazon (Berenguer et al. 2021) and can be used to synthesise the effects of fire on forest structure. However, future work would benefit from measuring the vertical distribution of the forest itself, through e.g., LiDAR. Wood density may be a weaker proxy of temporal changes in forest structure during post-disturbance recovery, and structural measures such as canopy height are known to be strong indicators elsewhere in the tropics (Santos et al. 2024). Structural proxies are also scalable, as they can be detected by satellites including GEDI (Dubayah et al. 2020) and the new BIOMASS mission (Quegan et al. 2019).

However, if microclimate buffering can be predicted by aboveground carbon stocks (or other measures of forest structure) across a wide range of disturbances intensities and recovery timelines, then this relationship can be used to predict thresholds when the forest can no longer buffer macroclimate. In our study, temperatures at the height of 0.5 m (which is the highest buffering during the day) varied in a predictable way, such that a forest loses 1.55 °C of buffering capacity for every 50 MgC/ha lost, and reached the same temperature as above the canopy (*i.e.*, zero buffering) with a carbon stock of 72.10 MgC/ha.

Our results from five vertical profiles showed strong changes in the vertical stratification of temperatures in Amazonian forest due to fire disturbance, and highlight that the third dimension of microclimate variability – height – interacts with the horizontal and temporal dimensions and hence requires further research attention. Future research should focus on replicating the vertical transects across a wider range of disturbances and times since disturbance, assessing variation across the calendar year; and expanding to other regions of the Amazon that have different macroclimatic regimes, for example with different dry season lengths (Carvalho et al. 2021). In addition, more accurate measurements of the vertical structure of forest would help to identify key drivers of change in microclimatic conditions from the canopy to the forest floor. For instance, 3D scanning using Terrestrial Laser Scanning (TLS) and airborne Light Detection and Ranging (LiDAR) can provide precise (sub-meter-scale) information on canopy height, leaf area index (LAI), gap fraction, and crown dimensions (Zellweger et al. 2019). This can improve the modelling and mapping of microclimate and allow large scale quantification of microclimate stratification across the Amazon. Finally, research on the biotic responses to changing microclimates would benefit from examining changes in the vertical profile as well as across space.

## ACKNOWLEDGEMENTS

This work was supported by the Natural Environment Research Council (NERC) RAINFAUNA project NE/X015262/1, multiple other NERC grants (NE/F01614X/1, NE/G000816/1, NE/K016431/1, NE/S01084X/1, and NE/X019039/1), PELD-RAS (CNPq, Brazil/CAPES, Brazil/PELD, Brazil 441659/2016-0), and the BNP Paribas Foundation, France, Climate and Biodiversity Initiative (Project Bioclimate). We thank the Large-Scale Biosphere-Atmosphere Program for logistical and infrastructure support during field measurements in Santarém. We are deeply grateful to Gilson Jesus Oliveira (Xarope), Renilson Menezes de Freitas (Graveto) and Jarilson Garcia Vilar for the help with fieldwork. This paper is number 135 in the Rede Amazônia Sustentável publication series https://ras-network.org/.

## SUPPLEMENTARY MATERIAL

**Figure S1.**
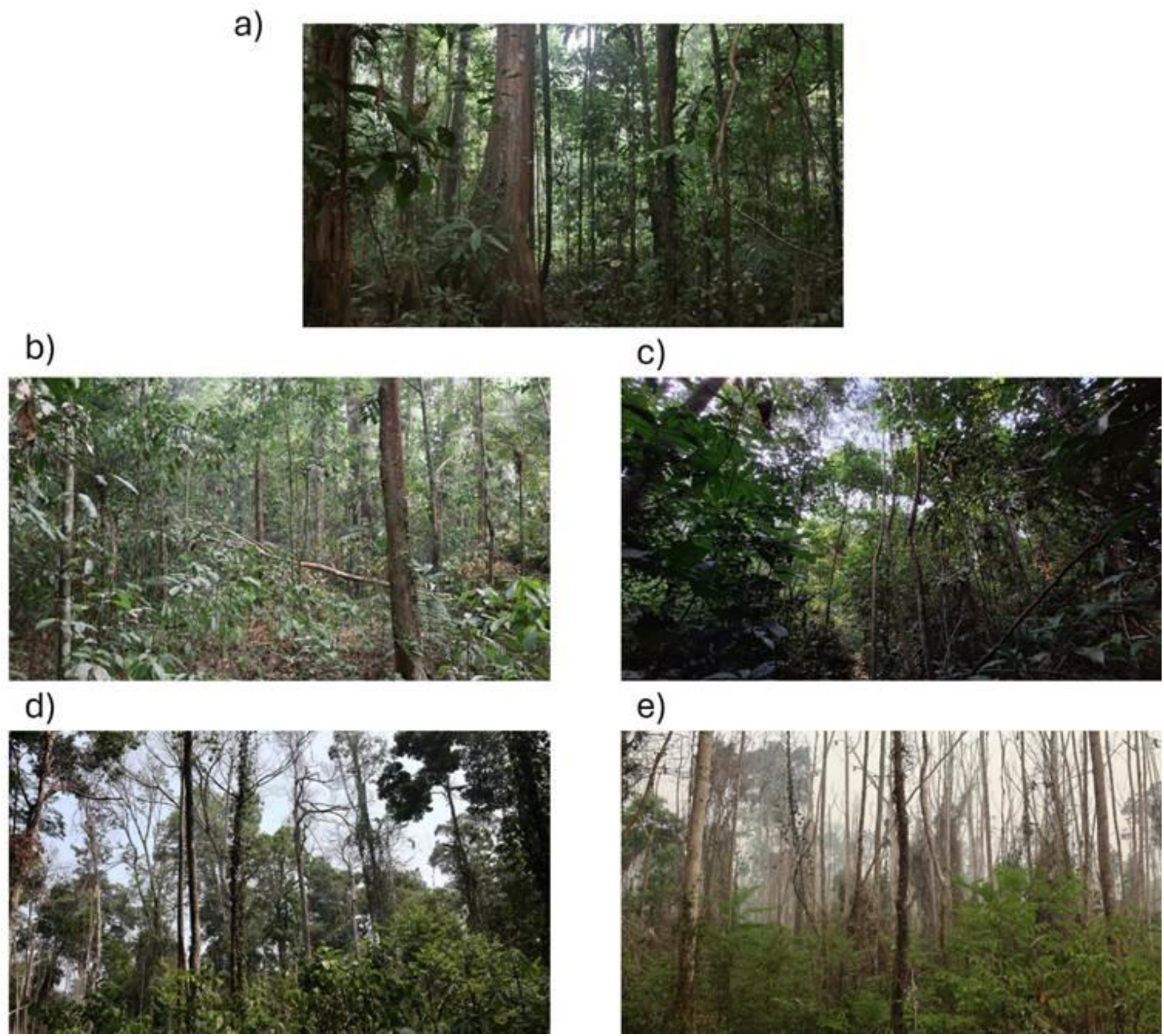
Amazonian forests along an anthropogenic disturbance gradient. a) Undisturbed primary forest with a closed canopy and a relatively open understorey. Forests in b) and c) are primary forests that burned once in 2015 during the extreme drought caused by an El Niño event. Forests in d) and e) are primary forests that burned twice, first in 2015 and second in 2023 during the extreme droughts caused by El Niño events. Photos were taken in November 2024 in Tapajós National Forest reserve (3°31’01” S, 55°04’23” W), Pará State, Brazil.

**Figure S2.**
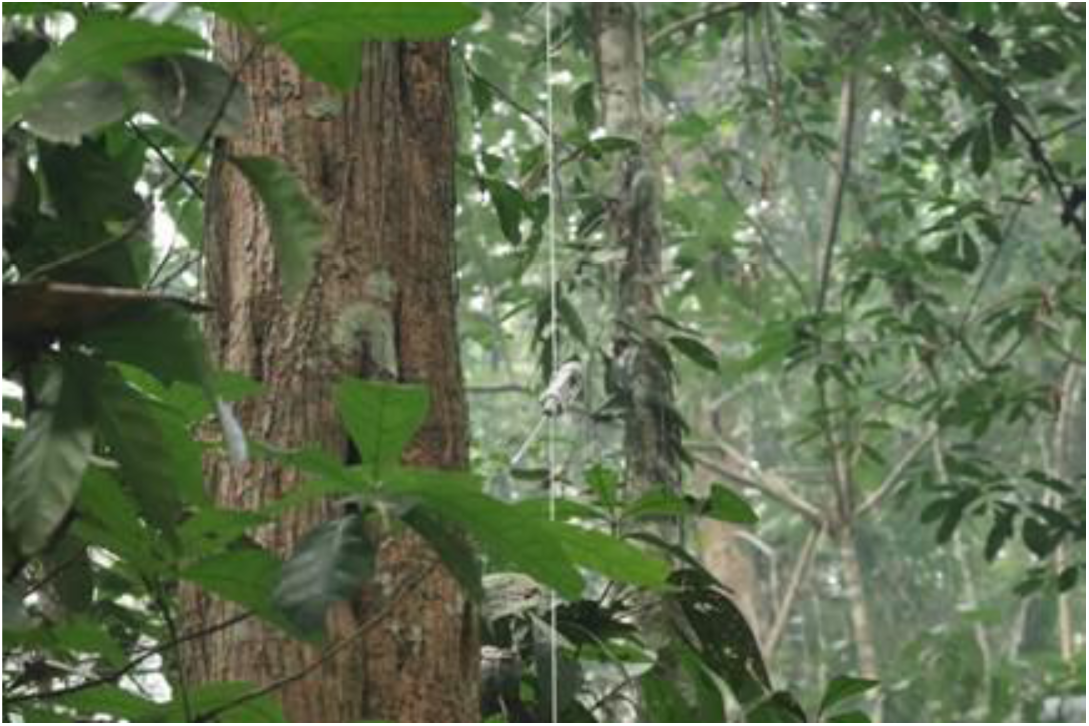
Temperature datalogger in the vertical transect. SurveyTag (https://surveytag.co.uk/) temperature dataloggers were attached to a nylon rope at known heights from the ground to measure temperatures along the vertical profile in undisturbed and disturbed forests in Tapajós National Forest reserve (3°31’01” S, 55°04’23” W), Pará State, Brazil.

## Notes

### Competing Interest Statement

The authors have declared no competing interest.

